# Tri-AD: Hippocampal cell-type-specific responses to age, sex and APOE genotype

**DOI:** 10.64898/2025.12.03.691944

**Authors:** Yaqiao Li, Yanxia Hao, Carlota Pereda Serras, Luokang Yao, Zherui Liang, Ana Almonte-Loya, Jessica Blumenfeld, Kaylie Suan, Seo Yeon Yoon, Brian Grone, Leonardo Ding, Yichuan Ma, Shiyao Sun, Boris Oskotsky, Tomiko Oskotsky, Carlo Condello, Yadong Huang, Marina Sirota

## Abstract

Alzheimer’s disease (AD) risk is strongly shaped by age, sex, and the apolipoprotein E ε4 (*APOE4*) allele—the strongest genetic risk factor for late-onset AD. While each factor has been studied independently, their combined impact on cellular and molecular processes remains unclear. Here, we used single-nucleus RNA sequencing (snRNA-seq) to profile hippocampal cell states in a sex-balanced cohort of human *APOE4/4* and *APOE3/3* knock-in mice across 6, 12, and 18 months of age. We identify sex as the major driver of variation in cell-type abundance and find that oligodendrocytes exhibit pronounced male-biased transcriptional sensitivity to APOE4. Differential expression and cell–cell communication analyses further reveal sex-divergent temporal trajectories in inhibitory neurons, with females showing early APOE4-associated suppression of synaptic pathways and males displaying a delayed but convergent decline. Together, these findings clarify how age, sex, and *APOE* genotype jointly regulate cell-type-specific gene expression and intercellular communication in the aging hippocampus, providing an innovative and publicly accessible database for aging and AD research and related precision medicine.

## Introduction

Over 55 million people worldwide are living with Alzheimer’s Disease (AD), and an estimated 416 million are affected by prodromal, preclinical, or AD dementia stages^1,2^. Despite decades of drug discovery efforts, available treatment options remain limited and often demonstrate variable efficacy across patient populations^3–5^. This is due partly to the substantial biological and clinical heterogeneity of AD, which hinders therapeutic development^6–8^. Thus, understanding the diverse pathogenic mechanisms that shape disease progression in different patient contexts is critical for precision medicine.

Among all risk factors for AD, growing evidence indicates that age, sex, and *APOE* genotype are significant non-modifiable contributors and critical focal points for advancing precision treatments^9,10^. Aging is the strongest risk factor, with AD prevalence rising sharply after age 65^11^. Sex differences further complicate disease manifestation^12,13^. Women make up approximately two-thirds of AD cases in the U.S.; a pattern that persists globally even after accounting for longer life expectancy in women^14^. The apolipoprotein E (*APOE*) *ε4* allele is the strongest genetic risk factor for late-onset AD, with over 50% of AD patients carrying at least one copy of the *APOE4* gene^15^. *APOE4* carriers typically exhibit exacerbated amyloid-β (Aβ) plaques and tau neurofibrillary tangles, the two pathological hallmarks of AD^16,17^. Notably, these major risk factors interact. Female *APOE4* carriers are at higher risk of developing AD at earlier ages, show greater tau pathology, and exhibit more pronounced hippocampal atrophy in mild cognitive impairment (MCI) than male carriers^18^. Together, these risk factors not only influence disease susceptibility but may also interact in complex ways that shape disease progression and therapeutic response.

A deeper understanding of AD molecular heterogeneity is essential for refining patient subtypes and developing targeted interventions^19,20^. Growing evidence indicates that APOE4 exerts heterogeneous, multicellular effects in AD^21–23^. A recent review proposed a cell-type-specific APOE4 cascade that initiates when stressed neurons upregulate APOE4, become dysfunctional, and release inflammatory signals that activate astrocyte and microglia, ultimately amplifying neuroinflammation and driving synaptic, neuronal, and myelin loss^24^. Whether this cascade generalizes across sexes, however, remains unclear. Human transcriptomic studies at advanced age consistently reveal marked glial pathway alterations associated with APOE4^25^ or female sex^26^, most prominently neuroinflammatory activation in astrocytes and microglia, consistent with late-stage features of the proposed cascade. However, demographic and sampling constraints inherent to human studies, such as the scarcity of *APOE4/4* donors, hinders systematic investigations of temporal dynamics and the early neuronal initiation phase^27,28^. Animal studies further support a sex-dimorphic APOE4-dependent effect: female microglia in an *App* mouse model activate at earlier ages than males^29^, and female mice expressing human *APOE4* and overexpressing *APP* exhibit reduced Aβ plaque compaction and microglia coverage of plaque compared to their male or *APOE3/3* counterparts^30^. Despite these insights, most sex-stratified studies have focused primarily on microglia, leaving the interactive effects of sex and APOE4 across diverse brain cell types largely uncharacterized.

To address this gap, we investigated the combined influence of age, sex, and *APOE* genotype on hippocampal cell states using single-nucleus RNA sequencing (snRNA-seq) in a sex-balanced cohort of human *APOE4/4* and *APOE3/3* knock-in mice across 6, 12, and 18 months of age. Our dataset revealed sex as the dominant contributing factor to variance in cell type abundance. Oligodendrocytes displayed pronounced male-biased transcriptional sensitivity to APOE4, including stronger activation of stress-response pathways. Through differential gene expression and cell-cell communication analyses, inhibitory neurons exhibited sex-divergent temporal trajectories of APOE4-related dysfunction, with females showing early suppression of synaptic and signaling pathways and males demonstrating a delayed but ultimately convergent decline. By delineating how age, sex, and *APOE* genotype collectively shape cell-type-specific gene expression and intercellular interactions, this study provides critical insight into AD molecular heterogeneity and lays a foundation for developing precision therapeutic strategies tailored to distinct patient subgroups. Most importantly, this comprehensive dataset is publicly accessible on a code-free platform (ucsftriad.org), providing researchers with an intuitive resource for independent exploration. This high-quality, fully sampled dataset enables researchers to investigate mechanistic hypotheses and explore biological questions that current human datasets cannot readily address due to their limited demographic or genetic representation.

## Results

### Cell-type-specific transcriptomic profiling of the mouse hippocampus across age, sex, and human *APOE* **genotypes**

To systematically investigate the specific and interactive transcriptomic effects of the three major risk factors associated with AD, we generated a comprehensive snRNA-seq dataset comprising hippocampal samples from twelve experimental conditions (n = 4 mice per condition, except n = 3 for 6 month *APOE3/3* males), defined by factorial combinations of age (6, 12 or 18 months), sex (male or female), and human homozygous *APOE* genotype (*APOE3/3* or *APOE4/4*) (Figure 1A). Focusing on six disease-relevant brain cell types—excitatory neurons (Ex.Neu), inhibitory neurons (In.Neu), astrocytes (Ast), microglia (Mic), oligodendrocytes (Oli), and oligodendrocyte precursor cells (OPCs)— we evaluated the effects of age, sex, and *APOE* genotype on cell-type abundance, differential gene expression, pathway enrichment, and intercellular communications across the twelve distinct experimental conditions (Fig. 1a).

**Fig. 1.**
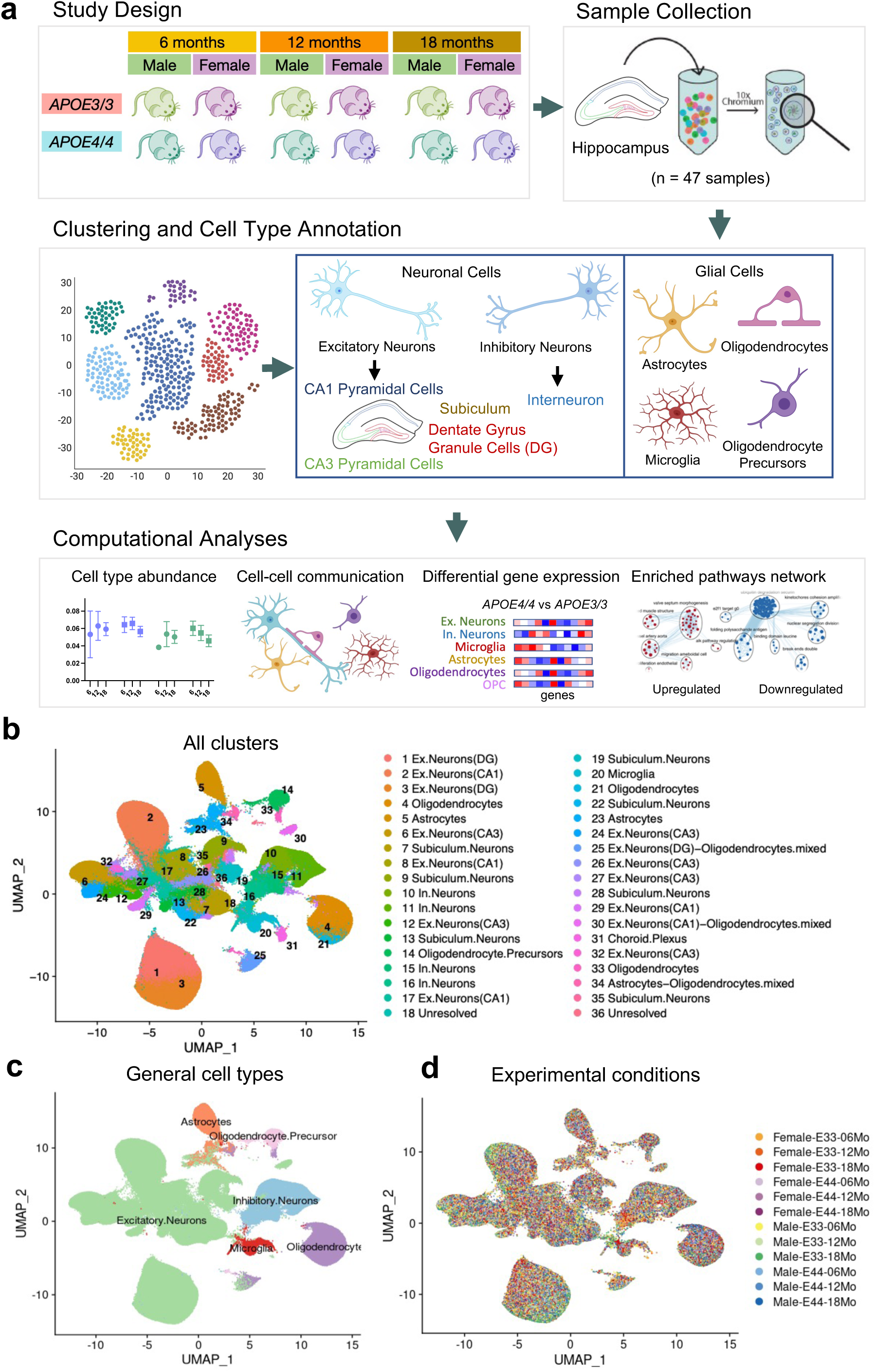
| Study overview. **a.** Overview of the study design, including cohort design, nuclei isolation workflow for snRNA-seq, and downstream computational analyses. **b.** UMAP embedding showing 36 transcriptionally distinct cellular populations, annotated by major cell types with resolved neuronal subtypes. **c.** UMAP annotated by the six major brain cell types. **d.** UMAP annotated by the twelve experimental conditions defined by varying combinations of the three risk factors—age, sex, and *APOE* genotype.

After standard processing and quality control, we generated a filtered dataset comprising 27,153 gene features across 555,008 nuclei for further analysis. Using graph-based clustering and Uniform Manifold Approximation and Projection (UMAP) for visualization, we identified 36 distinct cell clusters (Fig. 1b). By comparing their expression profiles to established marker gene sets (Extended Data Fig. 1a), these clusters were annotated into major brain cell types and the neuronal clusters were subcategorized into CA1 pyramidal cells (CA1), CA3 pyramidal cells (CA3), dentate gyrus granule cells (DG), subiculum neurons, and interneurons (Fig. 1b and Extended Data Fig. 1b). The clusters were well-defined and segregated into the six major cell types (Fig. 1c), with a uniform distribution across ages, sexes, and *APOE* genotypes (Extended Data Fig. 1b). UMAP visualization of all nuclei colored by experimental conditions shows that the twelve experimental conditions are broadly intermixed across clusters, with no condition-specific segregations (Fig. 1d). As expected, high levels of human *APOE* expression were visually apparent in astrocyte and microglia clusters, with moderate expression in neurons and oligodendrocytes, whereas mouse *Apoe* expression was negligible (Extended Data Fig. 1c).

### Sex is the strongest contributing factor to observed variance in cell type abundance

Cell type abundance was quantified as the proportion of nuclei from each cell type captured in each biological sample (Extended Data Fig. 2a). Three-way ANOVA was performed to assess the main and interaction effects of age, sex, and *APOE* genotype on the observed variations in cell type proportions across the twelve experimental conditions. This analysis revealed that sex most consistently emerged as a significant contributing factor to variance in cell-type abundance (Fig. 2a). Sex significantly influences proportional variance observed in astrocytes, dentate gyrus excitatory neurons, inhibitory neurons, and OPCs (Fig. 2a,b). Additionally, age, *APOE* genotype, and their interaction were also significant drivers of variation in OPC proportions (Fig. 2a). Specifically, *APOE4/4* mice exhibited higher OPC fractions compared to *APOE3/3* mice at 6 months of age, with this difference diminishing with aging, suggesting an interactive effect between age and *APOE* genotype (Fig. 2b). For microglia, both sex and age emerged as the main contributors to variance, although neither factor reached statistical significance (Fig. 2a and Extended Data Fig. 2b).

**Fig. 2.**
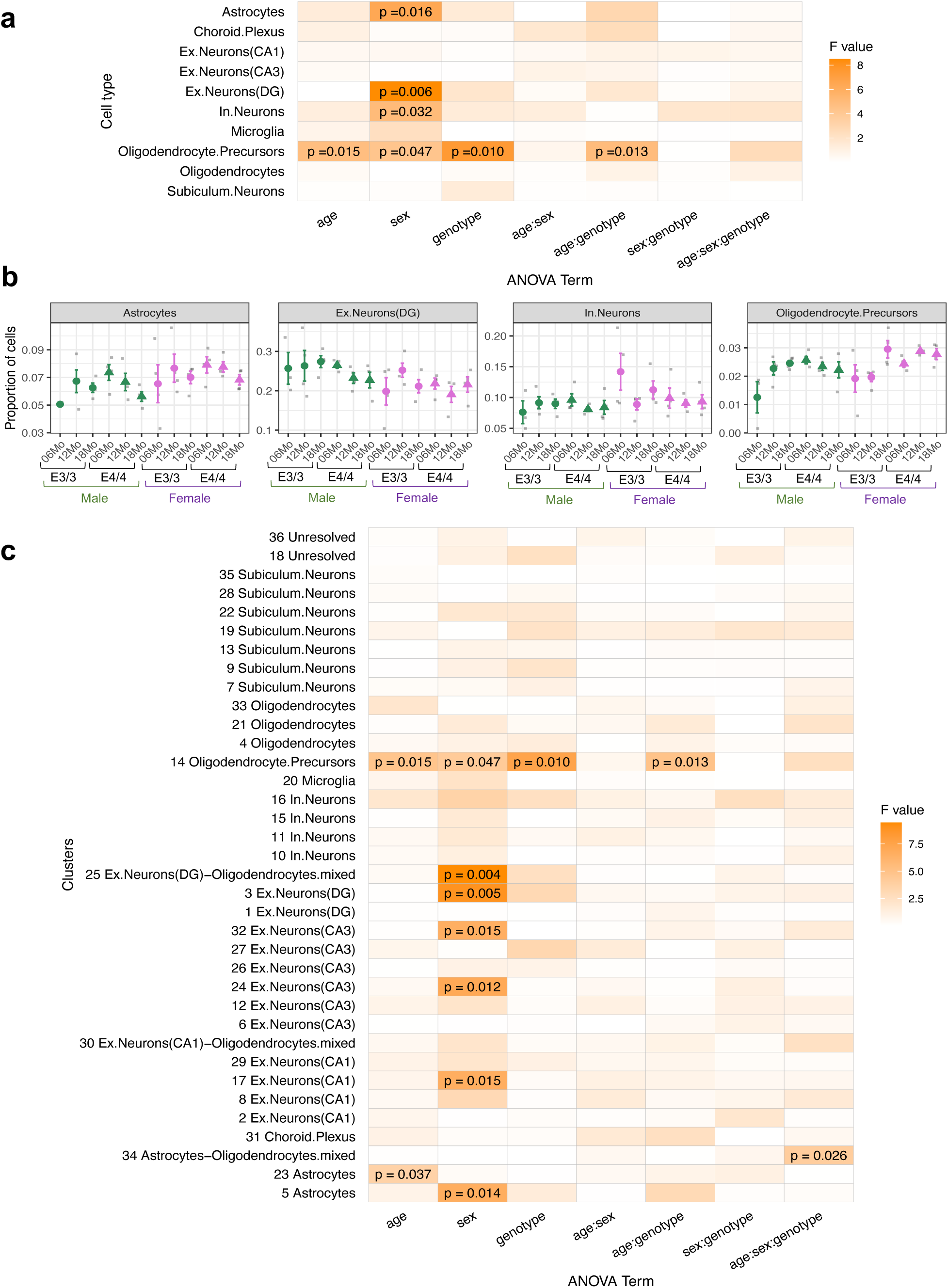
| Three-way ANOVA identifies sex as the dominant contributor to variance in cell abundance across conditions. **a.** Heatmap summarizing three-way ANOVA results for major cell type proportions. Color intensity reflects the magnitude of variance explained by each risk factor (age, sex, *APOE* genotype) and their interaction terms. P-values are shown for factors or interactions that significantly contribute to variance. **b.** Cell type proportions across the twelve experimental conditions for astrocytes, dentate gyrus excitatory neurons, inhibitory neurons, and OPCs, illustrating representative examples of factor-driven shifts in abundance. **c.** Heatmap of three-way ANOVA results for subcluster-level proportions. Color intensity denotes the extent of variance explained by each factor or their interaction, with significant effects labeled by their corresponding p-values.

Additionally, we further examined the impact of individual risk factors and their interactive effects on cell abundance across all 36 clusters. For each biological sample, cluster cell fractions were calculated as the number of cells in a given cluster divided by the total number of cells in that sample. Consistently, sex emerged as a significant factor contributing to the variance in cell abundance across several neuronal clusters (Fig. 2c). Notably, females exhibited higher fractions of CA1 pyramidal cells (cluster 17), while males showed higher abundances in the dentate gyrus (cluster 3) and CA3 pyramidal subpopulations (clusters 24 and 32) (Extended Data Fig. 2c). While astrocyte abundance collectively varied by sex, we observed a subtype (cluster 23) that consistently varied by age across sexes and genotypes (Fig. 2c and Extended Data Fig. 2c).

Taken together, all these data demonstrate that sex emerges as the most significant factor influencing cell type abundance, across both neurons (inhibitory neurons and dentate gyrus excitatory neurons) and glial cell populations (astrocytes and OPCs), potentially suggesting that sex-driven variations in cellular composition may underlie differential vulnerability to neurodegenerative processes. OPCs are further affected by age and genotype, which could suggest that these factors influence maturation of OPCs into oligodendrocytes. Furthermore, the sex dimorphism in neuronal subtype abundance elucidates sex-specific regional differences in mouse brains.

### *APOE4/4* drives sex-specific transcriptomic alterations across hippocampal cell types that manifest in age-dependent patterns

To systematically investigate transcriptomic alterations associated with age, sex, *APOE* genotype, and their interactions in the development of AD, we conducted pairwise differential gene expression analyses comparing *APOE4/4* (AD risk genotype) to *APOE3/3* (neutral/reference genotype) samples. By stratifying the analysis, we delineated how aging and sex modulate *APOE* genotype-related transcriptomic changes related to AD pathogenesis.

The resulting cell-type-specific differentially expressed genes (DEGs) revealed substantial variation in *APOE4*-dependent transcriptomic effects across ages and between sexes, with males overall exhibiting greater differences between *APOE4/4* and *APOE3/3* than females, reflected by consistently higher number of significant DEGs across most cell types and ages (Fig. 3a). However, notable exceptions were observed in inhibitory neurons and microglia at 18 months, where females displayed more significant DEGs, suggesting a stronger *APOE4/4-*associated influence in these cell types. Interestingly, inhibitory neurons in both sexes showed predominantly upregulated DEGs in earlier ages (6 and 12 months), whereas by 18 months, male exhibited mostly downregulated DEGs and female showed a mixture of up- and downregulated genes, revealing a sex-dimorphic transcriptomic response to APOE4 that becomes more pronounced in old age. Excitatory neurons followed a similar pattern, with predominately upregulated genes in young and adult mice but largely downregulated DEGs in males and a mixture response in females at 18 months. In glial cell types, DEG numbers generally peaked at middle age (12 months) and declined with aging in both sexes, except for microglia, which peaked at 18 months. Notably, male oligodendrocytes exhibited pronounced *APOE4/4-*associated transcriptomic alterations at all ages, whereas female oligodendrocytes showed a marked increase in DEGs at 12 months (Fig. 3a). Together, these findings highlight that *APOE4/4* drives temporally distinct transcriptomic effects in neurons and glial cells across sexes, with males showing heightened sensitivity to APOE4 across most cell types.

**Fig. 3.**
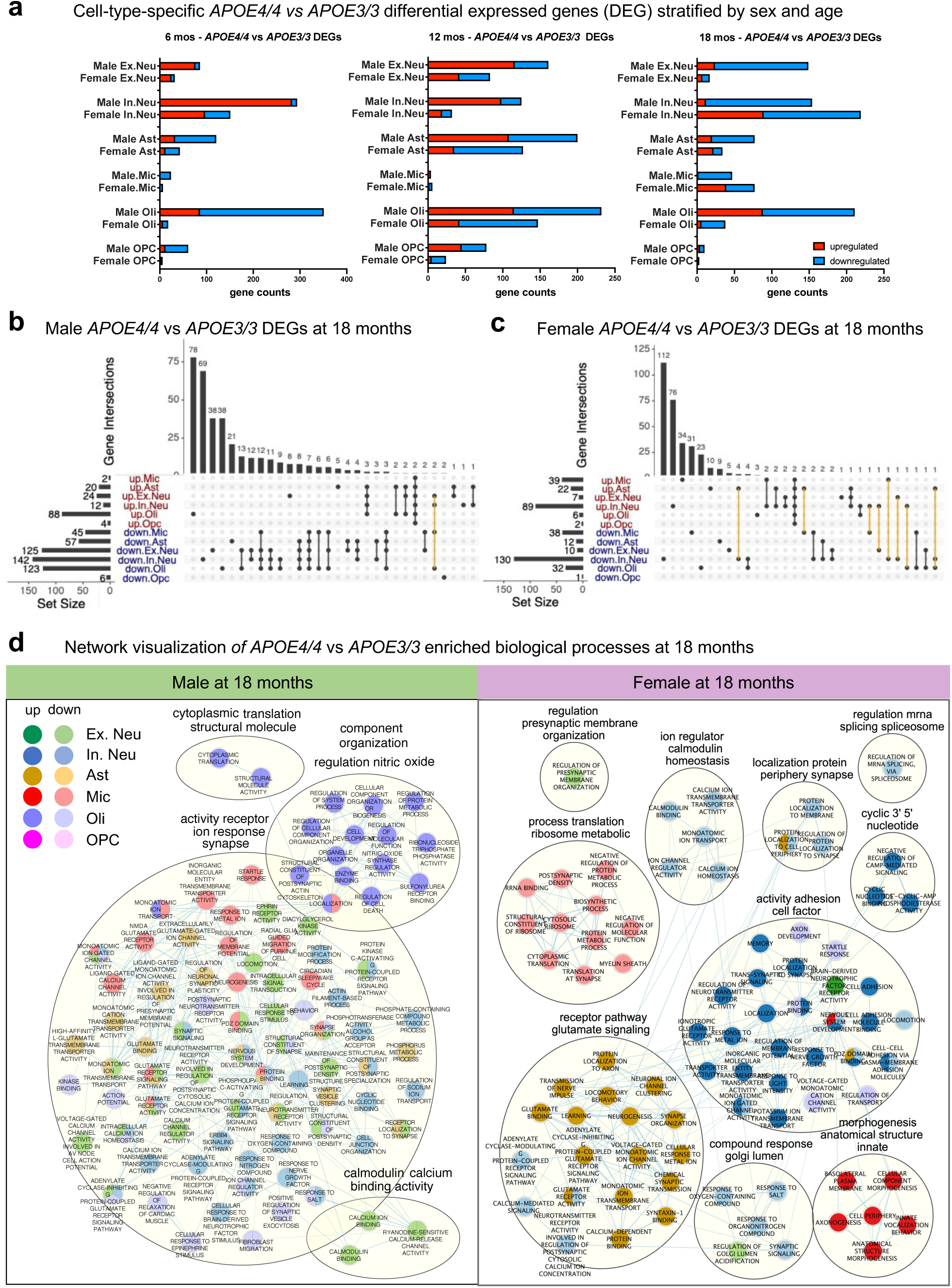
| Differential gene expression analysis of *APOE4/4* versus *APOE3/3*, stratified by sex and age. **a.** Cell-type-specific differentially expressed gene (DEG) counts for the six major cell types in males and females at 6, 12, and 18 months. Red bars indicate upregulated genes and blue bars indicate downregulated genes. **b,c.** UpSet plots showing DEG overlaps (*APOE4/4* vs. *APOE3/3*) at 18 months among the six major cell types in males (**b**) and females (**c**). Yellow connecting lines highlight DEG intersections across cell types that exhibit opposite directions of regulation. **d.** Enriched pathway networks for males and females at 18 months comparing *APOE4/4* mice to *APOE3/3* mice, stratified by cell type. Each node represents a significantly enriched pathway, grouped into biological themes and linked to functionally related clusters. Node color indicates the associated cell type(s), with darker hues representing upregulated pathways and lighter hues representing downregulated pathways.

### *APOE4/4* induces male-biased oligodendrocyte and female-biased inhibitory-neuron transcriptomic changes at advanced age

A cross-cell-type comparison of *APOE4/4* versus *APOE3/3* DEGs at 18 months (corresponding to late adulthood in humans) revealed that the majority of DEGs were unique to a single cell type in both males and females (Fig. 3b,c and Extended Data Fig. 3), indicating substantial cell-type-heterogeneity in APOE4-associated transcriptomic alterations. While a small subset of DEGs was shared across cell types, these overlapping DEGs were often regulated in opposite directions in females (Fig. 3c), suggesting stronger cell-type-divergent transcriptional responses to *APOE4/4* in females at old age. In contrast, nearly all overlapping DEGs in males were regulated in the same direction across cell types (Fig. 3b), indicative of more coordinated transcriptomic effects by *APOE4/4* across cell types.

Additionally, we investigated pathway enrichment by *APOE4/4* at 18 months and visualized the results as networks of biological clusters (Fig. 3d). In males, most *APOE4/4* versus *APOE3/3* pathways were enriched by downregulated DEGs linked to ion response, synapse, and receptor activity. Conversely, upregulated pathway enrichments were observed exclusively in oligodendrocytes, involving cytoplasmic translation, cellular organelle regulation, and nitric oxide activity. Notably, nitric oxide has been implicated in oligodendrocyte death and the suppression of myelin gene expression^31^, and APOE4 has been found to impair myelination in both human aging brains and *APOE4*-carrying mice^21^. Furthermore, evidence suggests that APOE4-associated myelination deficits are sex-dependent, with severe demyelination and a higher abundance of disease-associated oligodendrocytes (DAO) observed in male APOE4 mice^32^, findings consistent with our sex-specific transcriptomic signatures in oligodendrocytes.

In contrast to males, *APOE4/4*-enriched biological processes in females at 18 months were predominantly upregulated, including glutamate signaling pathways in astrocytes, cell adhesion processes in inhibitory neurons, and cellular morphogenesis in microglia, suggesting potential reactive or compensatory remodeling of glial and neuronal networks (Fig. 3d). At the same time, several critical pathways were downregulated, including glutamate signaling, calmodulin channel homeostasis, and protein localization in inhibitory neurons and protein biosynthetic processes in microglia. Together, these findings suggest that female *APOE4/4* brains exhibit a mixed transcriptional response, with upregulation of structural and signaling pathways alongside suppression of metabolic and synaptic programs, consistent with a pattern of compensatory activation followed by functional decline in neural and glial cells. Overall, these sex-specific patterns at advanced age reveal that, while *APOE4/4* drives broad transcriptomic downregulation in males with the notable exception of heightened oxidative stress responses in oligodendrocytes, females display a more heterogeneous profile with both activating and degenerative signatures, particularly in inhibitory neurons.

### Male oligodendrocytes exhibit heightened transcriptional sensitivity to *APOE* genotype changes

Consistent with pathway enrichment analyses at 18 months showing a male-specific impact of *APOE4/4* on oligodendrocyte function (Fig. 3d), cross-age DEG comparisons of oligodendrocyte revealed substantially greater transcriptomic alterations in males than females at all ages (Fig. 4a and Extended Data Fig. 4), underscoring a male-biased oligodendrocyte vulnerability to APOE4. Examination of the DEG identities revealed that stress response and cellular energy metabolism genes were consistently upregulated by *APOE4/4* in males but often downregulated in females (Fig. 4b), suggesting a sustained state of metabolic stress and compensatory activation in *APOE4/4* male oligodendrocytes. Interestingly, genes related to synaptic vesicle release, ion channel excitability, postsynaptic plasticity, and neuronal morphogenesis were initially upregulated in *APOE4/4* male oligodendrocytes at 6 months compared to *APOE3/3*, but gradually declined with age, becoming significantly downregulated by 18 months (Fig. 4c). Although these functions are typically neuronal, their enrichment in oligodendrocytes likely reflects altered signaling between neuron and oligodendrocyte or aberrant activation of neuronal gene programs within oligodendrocytes^32,33^. This fluctuation, potentially driven by the transiently elevated *APOE* expression observed in *APOE4/4* males at 6 months (Extended Data Fig. 5), suggests an early compensatory attempt by oligodendrocytes to support neuronal activity that ultimately fails with aging, which could contribute to a progressive decline in synaptic support and myelination.

**Fig. 4.**
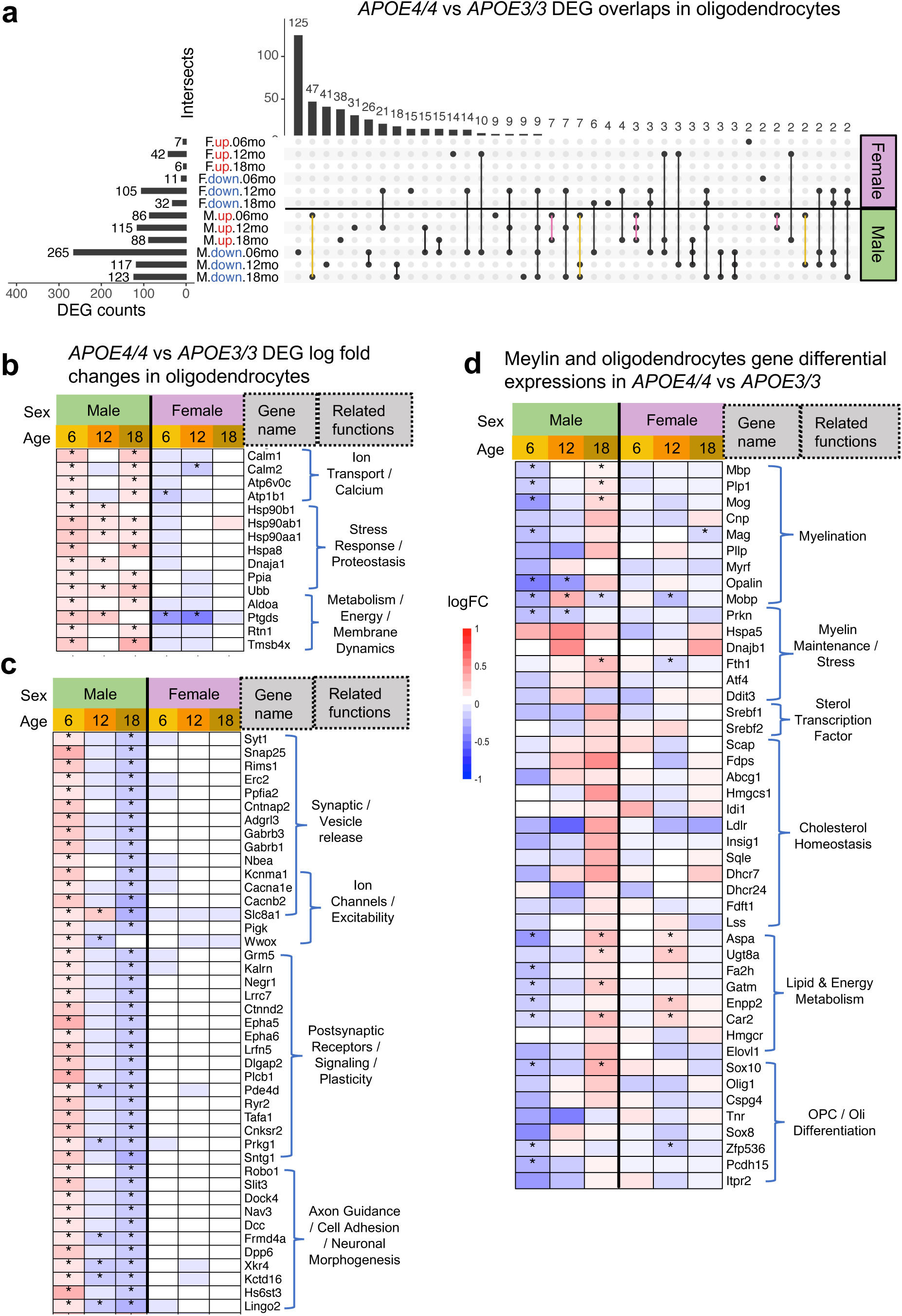
| APOE4/4 induces heightened oligodendrocyte transcriptional responses in males. **a.** UpSet plots showing oligodendrocyte DEGs (*APOE4/4* vs. *APOE3/3*) across sexes and ages, with each condition further split by direction of regulation. Pink lines highlight DEG intersections that are consistently upregulated in males with *APOE4/4* across all ages. The yellow lines highlight DEG intersections that are upregulated at 6 months but downregulated at 12 and/or 18 months. **b.** Heatmap showing expression patterns of male oligodendrocyte DEGs that are consistently upregulated with *APOE4/4* (highlighted by pink lines in **a**). **c.** Heatmap showing expression patterns of male oligodendrocyte DEGs that are upregulated at 6 months and subsequently downregulated with aging (highlighted by the yellow lines in **a**). **d.** Heatmap showing expression patterns of oligodendrocyte-relevant genes across sexes and ages. **b–d.** Colors indicate the direction and magnitude of regulation, with red denoting upregulation and blue denoting downregulation; significant differential expression is marked by an asterisk (*).

Since oligodendrocytes play a central role in myelin production and maintenance of neuronal myelin sheaths, we examined the differential expression of myelination-related genes in oligodendrocytes across sexes and ages (Fig. 4d). Previous studies in humans have shown that *APOE4* carriers exhibit downregulation of myelination genes accompanied by upregulation of cholesterol homeostasis genes^21^. Consistently, our results revealed upregulation of genes involved in lipid metabolism and cholesterol homeostasis, along with *APOE4*-associated suppression of myelin genes in both sexes across most ages, except in males at 18 months, where several myelin-associated genes (*Mbp, Plp1, and Mog*) were upregulated (Fig. 4d). While myelination genes were significantly downregulated at 6 months in males, this late upregulation may reflect a reactive or compensatory response to progressive myelin dysregulation, potentially driven by chronic stress signaling, altered lipid metabolism, or recruitment of OPCs attempting to restore myelin integrity. Overall, the data reveal a male-biased, *APOE4/4*-driven disruption of oligodendrocyte function, marked by early compensatory activation and gradual decline in neuronal support with aging.

### Early female and delayed male vulnerability in *APOE4/4*-associated inhibitory neuron dysfunction

Inhibitory neuron dysfunction has been increasingly recognized as an early driver of AD pathogenesis^24,28,34,35^. In our dataset, we observed significant *APOE4/4*-associated enrichment of pathways in inhibitory neurons at 18 months (Fig. 3d), indicating pronounced genotype-driven alterations in this vulnerable neuronal population. To further dissect these effects, we examined age- and sex-dependent transcriptomic responses to the *APOE4/4* genotype in inhibitory neurons (Fig. 5).

**Fig. 5.**
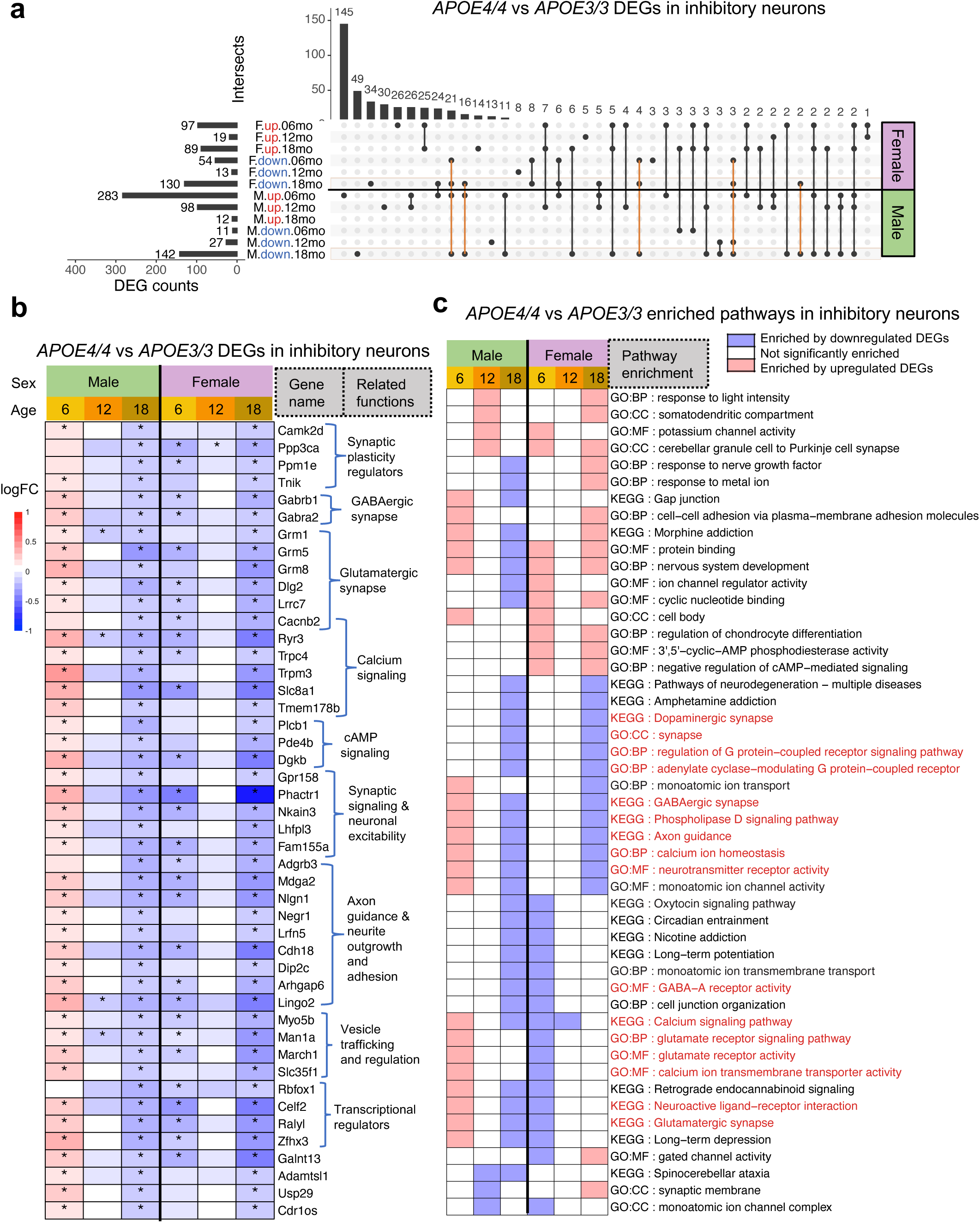
| Sex-divergent early responses converge with aging in inhibitory neuron dysfunction in *APOE4/4* mice. **a.** UpSet plots showing inhibitory neuron DEGs (*APOE4/4* vs. *APOE3/3*) across sexes and ages, with each condition further separated by direction of regulation. Orange lines highlight DEGs shared between males and females at 18 months. **b.** Heatmap displaying expression patterns of DEGs shared between females and males at 18 months (highlighted by orange lines in **a**). Colors indicate the direction and magnitude of regulation, with red denoting upregulation and blue denoting downregulation; significant differential expression is marked by an asterisk (*). **c.** Heatmap of *APOE4/4* versus *APOE3/3* enriched pathways in inhibitory neurons, stratified by age and sex. Colors indicate the direction of regulation, with red denoting upregulation, blue denoting downregulation, and white indicating non-significant changes in the corresponding groups.

At 18 months, all overlapping DEGs between males and females were downregulated by the *APOE4/4* genotype, with no shared upregulated DEGs (Fig. 5a), suggesting a sex-convergent decline in inhibitory neuron activity at advanced age. These genes were enriched for pathways related to synaptic transmission and neuronal signaling, including GABAergic and glutamatergic synapses, calcium and cAMP signaling, synaptic plasticity, and vesicle trafficking (Fig. 5b,c). Interestingly, females showed early downregulation of these genes at 6 months, persisting in some cases at 12 months, whereas males exhibited initial upregulation at 6 months, followed by downregulation beginning around 12 months (Fig. 5b). This sex-specific trajectory suggests that male inhibitory neurons may undergo transient compensatory activation of synaptic and signaling pathways in response to *APOE4/4*-associated stress at 6 months, consistent with the elevated *APOE* expression observed in *APOE4/4* males (Extended Data Fig. 5). In contrast, females display early transcriptional suppression of these pathways, potentially contributing to premature synaptic and network dysfunction. The later decline in males likely reflects a loss of compensatory capacity over time, culminating in a shared inhibitory signaling deficit by 18 months in both sexes.

### Early female- and *APOE4*-dependent decline in cell–cell communication with aging

To assess whether the synaptic decline observed in inhibitory neurons extends to other cell populations, we examined synaptic transmission and intercellular signaling using cell–cell communication (CCC) analysis across all six cell types (Fig. 6). CCC analysis infers ligand–receptor interactions between cell types based on the transcriptomic data, quantifying the strength and number of signaling connections, with directional granularity distinguishing sender and receiver cell populations. To delineate the differential aging trajectories of intercellular communication specific to each sex and genotype background, we performed comparison across ages rather than between *APOE* genotypes.

**Fig. 6.**
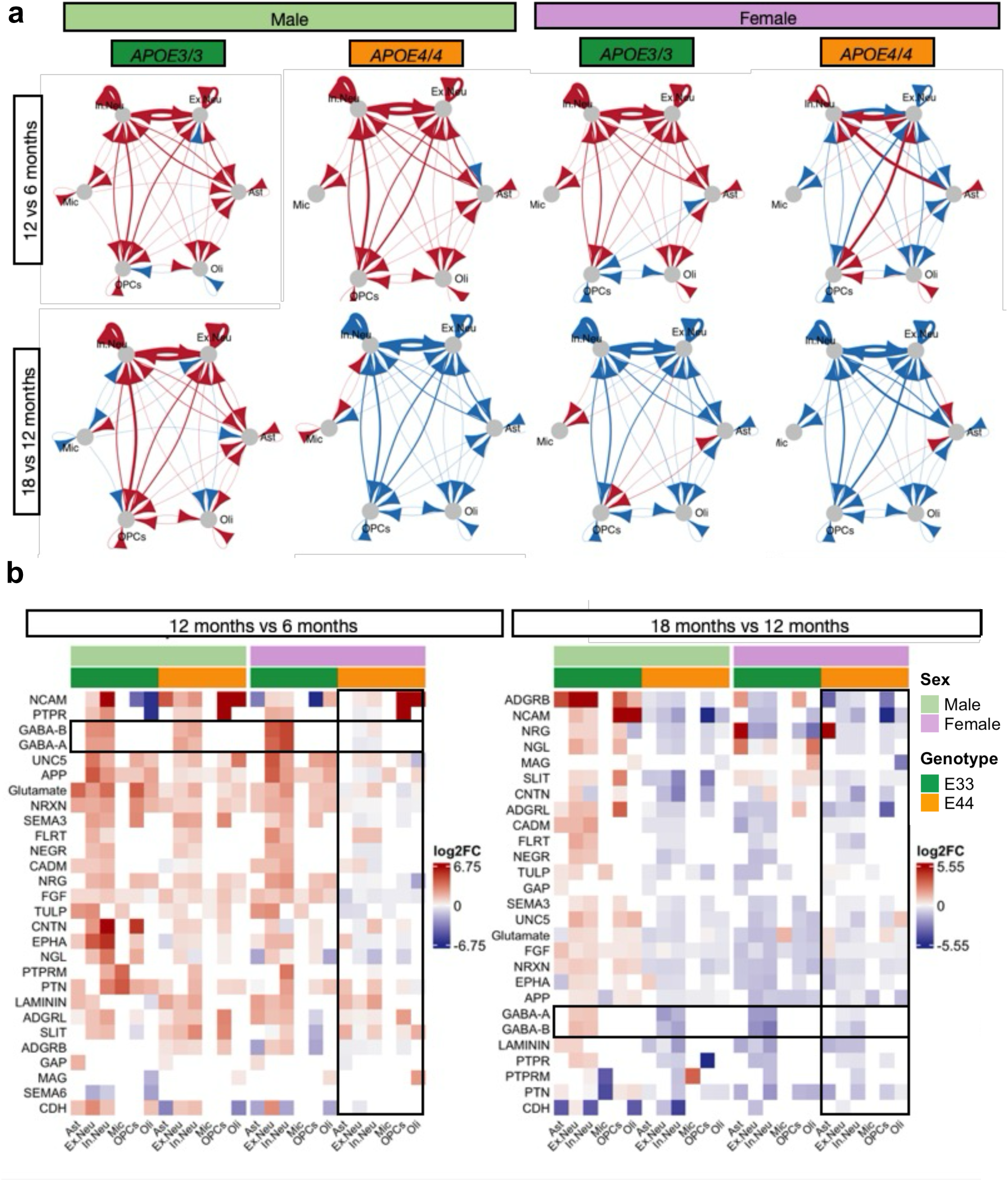
| Early declines in cell–cell communication with aging are driven by female sex and APOE4 genotype. **a.** Cell type–to–cell type communication strength inferred from ligand–receptor pair expressions, visualized as directed arrows from sender to receiver cell types. Red arrows indicate increased communication in older versus younger mice (12 vs. 6 months or 18 vs. 12 months), whereas blue arrows indicate decreased communication. **b.** Heatmap showing pathway-level CCC differences between older and younger mice across *APOE* genotypes and sexes, stratified by cell type. Values represent the scaled CCC probability differences between age groups. Signaling pathways correspond to ligand–receptor pairs mapped to KEGG pathway categories.

Pairwise age comparisons, 12 vs. 6 months (early aging) and 18 vs. 12 months (late aging), for male and female *APOE3/3* and *APOE4/4* mice revealed sex- and age-dependent alterations in CCC (Fig. 6a). During early aging, overall communication strength increased with age under *APOE3/3* conditions in both sexes, potentially reflecting normal network maturation. In contrast, female *APOE4/4* mice exhibited an early and pronounced reduction in signaling activity by 12 months, especially by decreased outgoing signaling from inhibitory neurons to other cell types, indicative of a premature loss of inhibitory network coordination. Interestingly, incoming signaling to inhibitory neurons from excitatory neurons, astrocytes, and oligodendrocytes was elevated at 12 months, suggesting a compensatory response by other cell types attempting to restore the disrupted inhibitory communication.

By 18 months, male *APOE4/4* and female *APOE3/3* mice exhibited a comparable global decline across nearly all cell types, mirroring the earlier changes observed in *APOE4/4* females (Fig. 6a). At this late stage, both sexes displayed markedly weakened intercellular signaling, with strongest reduction in CCC observed in *APOE4/4* mice and the mildest decline in *APOE3/3* males, suggesting a widespread communication breakdown associated with aging in females and the APOE4/4 genotype. Together, these findings suggest that females undergo an earlier disruption of CCC, especially in relation to inhibitory neurons, that is exacerbated by *APOE4/4*, whereas males experience a delayed but ultimately convergent decline, consistent with the sex-specific temporal vulnerability and inhibitory neuron signaling deficits observed in the DEG analysis.

When decomposing the CCC network into individual ligand-receptor pairs, we found that most synapse-related signaling pathways were moderately reduced in *APOE4/4* females during early aging, particularly GABA-A and GABA-B interactions among neurons (Fig. 6b and Extended Data Fig. 6). By 18 months, all groups except *APOE3/3* males exhibited further reductions in GABA-A and GABA-B signaling, supporting a progressive loss of inhibitory synaptic communication with aging that occurs earlier and more severely in *APOE4/4* females. Together, these findings suggest that *APOE4/4* and female sex jointly accelerate the breakdown of GABAergic communication and loss of inhibitory signaling with aging.

## Discussion

While aging, female sex, and the *APOE4* genotype represent the strongest risk factors for AD, their multifaceted interactions and combined influence on cellular states and molecular pathways remain elusive^9,10^, posing a major challenge to mechanistic understanding and therapeutic development. Systematic characterization of human molecular changes across age, sex, and *APOE* genotype is hindered by the limited sample availability of key patient subgroups, such as *APOE4/4* carriers^27,28^. Here, we present a comprehensive single-cell transcriptomic dataset characterizing the effects of age, sex, and *APOE4* genotype on hippocampal cell type composition, transcriptional profiles, and intercellular relationships in a mouse model of APOE4-related late-onset AD. Our analysis highlights sex as a major determinant of variation in cell-type abundance, *APOE4*-driven differential gene expressions, and the progressive decline in cell-cell signaling with aging. To promote open discovery, this comprehensive dataset is made publicly available for intuitive, code-free exploration at ucsftriad.org.

Sex differences in cell type abundance have been documented in healthy brains across species^36,37^ and in aging brains^37–39^. In line with these observations, our data indicated that, compared with the effects of age or *APOE* genotype, sex exerts a stronger influence on cell-type abundance, particularly in inhibitory neurons, a subtype of excitatory neurons (DG), astrocytes, and OPCs, whereas age and *APOE* genotype effects were limited to OPCs. Subclustering of neuronal subtypes further revealed sex-dimorphic cell proportions in dentate gyrus, CA1 and CA3 pyramidal neurons. Together, these findings highlight the importance of incorporating sex as a critical biological variable when quantifying cell composition changes associated with disease, ensuring accurate interpretation of neurobiological and disease-related changes. Future work examining how these sex-dependent changes in cellular composition influence brain function may provide insight into the elevated AD susceptibility in females.

The selective vulnerability of inhibitory neurons has been consistently reported in AD patients and animal models^19,28,35,40–43^. Our *APOE4/4*-differential transcriptomic profiling stratified by age and sex revealed a temporal divergence in *APOE4/4*-associated inhibitory neuron dysfunction between males and females. Although inhibitory neuron abundance was generally higher in females across ages, this difference was most pronounced in APOE3/3 mice and substantially diminished in APOE4/4 mice (Fig. 2b), indicating that APOE4 erodes the female advantage in inhibitory neuron abundance. Consistent with this, females showed early-onset *APOE4/4*-driven suppression of synaptic and signaling pathways, indicating premature functional decline. In contrast, males exhibited a delayed trajectory, with initial upregulation of synaptic, calcium, and cAMP signaling pathways at 6 months, followed by downregulation beginning around 12 months and converging with the female decline by 18 months. This pattern suggests that female inhibitory neurons may be intrinsically more susceptible to *APOE4*-related stress, leading to earlier disruption of inhibitory signaling and network stability. Male inhibitory neurons, by contrast, appear to have transiently active compensatory mechanisms, reflected by the early pathway upregulations and higher human *APOE* expression in *APOE4/4* than *APOE3/3* only in 6-month males. These upregulations might be an attempt to preserve inhibitory function but ultimately deteriorate with aging. It is possible that females also go through this compensatory phase, earlier than 6 months of age, but was not captured by this dataset. By 18 months, both sexes converge toward a shared state of synaptic downregulation and reduced inhibitory signaling, indicating that *APOE4* might disrupt inhibitory neuron functions through sex-specific temporal trajectories that ultimately lead to a common late-stage deficit.

The complementary CCC analyses reinforce this model, demonstrating that *APOE4/4* females experience an early and more pronounced suppression of signaling network across hippocampal cell types, especially the loss of inhibitory neuron output and diminished GABAergic signaling, whereas males exhibit a delayed but comparable decline during advanced aging. These results highlight sex as a major modulatory factor that amplifies disease-driven neuronal vulnerability and network instability, shaping both the onset and trajectory of inhibitory and intercellular signaling deterioration. The earlier collapse of inhibitory communication in females provides a potential cellular and molecular basis for the greater prevalence and earlier onset of AD in women, which is further exacerbated by *APOE4* genotype. More broadly, these findings emphasize the importance of investigating sex-dimorphic biological factors, such as hormonal regulation, metabolic differences, and distinct transcriptional responses, in studies of neuronal aging and disease, as well as in development of therapeutic strategies aimed at stabilizing inhibitory network function and preserving intercellular signaling across the aging brain.

Previous study has shown that sex differences in myelin vulnerability vary with disease context^32^. In aging mice treated with a demyelinating agent, females exhibited more severe myelin loss and greater susceptibility to demyelination^32^. In contrast, in APOE4-driven tauopathy models, males showed stronger interferon responses, greater activation of disease-associated microglia, and demyelination^32^. Our data also demonstrate a male-biased vulnerability of oligodendrocytes to APOE4-driven pathology. Across all ages, *APOE4/4* male oligodendrocytes showed markedly greater transcriptomic alterations than females, with persistent upregulation of stress-response and metabolic pathways and an early, transient activation of synaptic and signaling programs in oligodendrocytes. This compensatory signature, likely driven in part by elevated *APOE* expression specifically in 6-month *APOE4/4* males, ultimately deteriorated with age, resulting in pronounced downregulation of neuronal support pathways by 18 months.

Another study revealed that APOE4 disrupts cholesterol homeostasis, leading to cholesterol accumulations in oligodendrocytes, reduced myelination, and impaired insulation and support of neuronal electrical activity^21^. In our dataset, although both sexes exhibited APOE4-associated downregulation of myelination genes and upregulation of cholesterol homeostasis pathways, males showed markedly stronger transcriptional responses in these key pathways. These patterns are consistent with prior reports yet demonstrate a sex-biased APOE4 effect that imposes greater functional stress on male oligodendrocytes. It remains possible that female oligodendrocytes experience substantial baseline stress, independent of *APOE4/4* genotype, such that additional APOE4-driven perturbation is less discernible, an interpretation that will require future studies to evaluate directly.

There are limitations in this study. First, this study focuses on investigating *APOE4* biological effects prior to pathology presence that could render the brain vulnerable to neurodegeneration. In the future, a mouse model that includes disease pathology, in addition to human *APOE4,* could further elucidate how *APOE4* and sex play a role in AD. Furthermore, novel omics technologies, such as spatial transcriptomics, will allow for a detailed study of how cell-cell interactions could lead to an increased vulnerability of female *APOE4* carriers, and will allow for the inclusion of extra-nuclear RNA Lastly, a thorough comparison and integration of mouse models with human data will be essential to assess the translatability of biological mechanisms across the two species and to determine what female APOE4-driven signatures observed in this study are conserved in human AD brains.

In summary, this study provides a comprehensive single-cell atlas that disentangles the intersecting effects of age, sex, and APOE4 genotype, three major risk factors whose combined impact on the aging brain has remained poorly understood. By revealing sex as a dominant modulator of APOE4-driven changes in inhibitory neurons, oligodendrocytes, and network-level signaling, our findings highlight distinct sex-biased temporal trajectories of vulnerability that ultimately converge on shared late-stage deficits. The discovery of sex-biased compensatory responses, differential susceptibility to APOE4-associated stress, and divergent patterns of synaptic, metabolic, and myelination-related dysfunction underscores the importance of incorporating sex and age as essential biological variables in mechanistic and translational AD research. Importantly, this publicly accessible dataset and interactive portal we provide (ucsftriad.org) offer a powerful, community-facing resource for investigators to interrogate how these three risk factors shape cellular phenotypes and molecular pathways in neurodegeneration. By enabling systematic, data-driven exploration of age–sex–genotype interactions, this work lays a foundation and provides a rich resource for future studies aimed at elucidating disease mechanisms and advancing precision therapeutic strategies for AD and other neurodegenerative disorders.

## Acknowledgements

This study was supported by the National Institute on Aging grants R01AG060393 to MS and R01AG057683 to MS and YH, RF1AG076647, R01AG078164, and P01AG073082 to YH, and NSF 2034836 to YL. We thank Sirota and Huang lab members for their valuable discussions about the experimental design as well as data analyses and interpretation. We also thank Eric Chow and the staff at the UCSF Center for Advanced Technology Core for advice and support with snRNA sequencing.

## Materials and Code Availability

All code used in this study is available in [GITHUB LINK]. Data is available for exploration and download in ucsftriad.org.

## Author contributions

YHuang and MS designed and supervised the study. YL and CPS performed the data analyses. YHao generated the animal cohort and isolated cell nuclei. YHao and KS prepared samples for snRNAseq. YL and LY created the Rshiny app at ucsftriad.org. ZL, AA, JB, SYY, BG, LD, YM, SS, BO assisted data collection and analysis. TO and CC provided advice and guidance on the study. YL, CPS, YHuang and MS wrote the manuscript. All authors have read and approved the manuscript.

## Methods Mice

All protocols and procedures followed the guidelines of the Laboratory Animal Resource Center at the University of California, San Francisco (UCSF). All mice had identical housing conditions from birth through sacrifice (12 h light/dark cycle at 19–23 °C and 30–70% humidity, housed five animals per cage, PicoLab Rodent Diet 20). ApoE3/3 and APOE4/4 homozygous knock-in mice^44,45^ (Taconic) were born and aged under normal conditions at the Gladstone Institute/UCSF animal facility. Both male and female APOE4/4 and APOE3/3 mice at 6, 12, and 18 months of age (n=4 mice per genotype per sex and per age, except n = 3 for 6 month *APOE3/3* males) were used for the study.

## cDNA library preparation and sequencing

Single-nuclei preparation for 10x loading. One frozen mouse hippocampus was placed into a pre-chilled 2 mL Dounce with 1 mL of cold 1X Homogenization Buffer (1X HB) (250 mM Sucrose, 25 mM KCL, 5 mM MgCl2, 20 mM Tricine-KOH pH7.8, 1 mM DTT, 0.5 mM Spermidine, 0.15 mM Spermine, 0.3% NP40, 0.2 units/µL RNase inhibitor, ∼0.07 tabs/ml cOmplete Protease inhibitor). Dounce with “A” loose pestle (∼10 strokes) and then with “B” tight pestle (∼10 strokes). The homogenate was filtered using a 70 µM Flowmi strainer (Bel-Art) and transferred to a pre-chilled 2 mL LoBind tube (Fischer Scientific). Nuclei were pelleted by spinning for 5 min at 4°C at 350 RCF. The supernatant was removed and the nuclei were resuspended in 400 µL 1X HB. Next, 400 µL of 50% Iodixanol solution was added to the nuclei and then slowly layered with 600 µL of 30% Iodixanol solution under the 25% mixture, then layered with 600 µL of 40% Iodixanol solution under the 30% mixture. The nuclei were then spun for 20 min at 4°C at 3,000 RCF in a pre-chilled swinging bucket centrifuge. 200 µL of the nuclei band at the 30%-40% interface was collected and transferred to a fresh tube. Then, 800 µL of 2.5% BSA in PBS plus 0.2 units/µL of RNase inhibitor was added to the nuclei and then were spun for 10 min at 500 RCF at 4°C. The nuclei were resuspended with 2.5% BSA in PBS plus 0.2 units/µL RNase inhibitor to reach at least 500 nuclei/µL. The nuclei were then filtered with a 40 µM Flowmi cell strainer. The nuclei were counted and then ∼13,000 nuclei per sample were loaded onto 10x Genomics Next GEM chip M. The snRNA-seq libraries were prepared using the Chromium Next GEM Single Cell 3ʹ HT kit v3.1 (10x Genomics) according to the manufacturer’s instructions. Libraries were sequenced on an Illumina NovaSeq 6000 sequencer at the UCSF CAT sequencing core.

## Sequence alignment, filtering, and counting

The demultiplexed fastq files were aligned to a custom reference genome, built from mm10-1.2.0 which includes introns, using the *cellranger count* function (version 4.0.0) with default parameters, as detailed in the Cell Ranger documentation. Subsequently, a single UMI count file per animal/sample was generated by the *cellranger count* function. Individual UMI count files were then combined into a single count matrix using *merge* function in Seurat package v4.0.4. Metadata, including age, sex, and genotype information, were added to each cell.

## Pre-processing and quality control

The count matrix was further processed with Seurat package^46^ by first calculating the percentage of mitochondria genes mapped per cell. The distribution of feature count, total mapped gene count, and percentage of mitochondria genes were visualized across biological samples as violin plots, and no obvious outlier was identified. We filtered the count matrix to only include cells with higher than 250 gene features, at least 500 gene counts, and mitochondria gene percentage lower than 1%. Potential misaligned or ambiguous gene features that expressed in fewer than 10 cells were also removed. These quality assurance steps resulted in a final Seurat object containing 27,153 gene features expressed by 536,498 nuclei.

## Normalization, dimensional reduction, and clustering

Count normalization and dimensionality reduction was conducted following standard procedure in the Seurat package^46^. In brief, we performed normalization and variance stabilization with an updated version of *sctransform*^47^, v2, and principal component analysis (PCA) with *RunPCA* (npcs = 30). Dimensional reduction through Uniform Manifold Approximation and Projection (UMAP) was performed with the *RunUMAP* function and considering the top 15 dimensions selected from the corresponding PCA.

The clustering was based on the first 15 principal components (PCs) using the *FindNeighbors* function in Seurat. This function embeds cells in a K-nearest neighbor graph, considering the Euclidean distance in PCA space and refining the edge weights between any two cells based on the shared overlap in their local neighborhoods. The clustering was obtained using the *FindClusters* function, which employs modularity optimization techniques such as the Louvain algorithm, with a resolution parameter of 0.5 and resulted in a set of 36 distinct clusters.

## Cell-type annotation

Major cell types, including astrocytes, microglia, oligodendrocytes, oligodendrocyte precursor cells, excitatory and inhibitory neurons, were classified using mouse brain cell markers in PanglaoDB^48^, a publicly available marker gene database. Further subdivision of hippocampus cell types, such as CA1 and CA3 pyramidal cells, were queried against hippocampal cell-type-specific marker genes as published in hipposeq^49^ (https://hipposeq.janelia.org). Cell type identities per cluster were determined by applying Seurat’s *AddModuleScore* function to sets of mouse brain marker genes. A module score for each cell type considered was calculated per cell. Each cell was assigned the corresponding cell type identity that generated the highest scores among scores for all cell types. If the highest and second highest scores of a cell were within 20% of the highest score, then the cells were deemed hybrids and excluded from further analysis. We assessed the validity of the assigned cell type identities by examining the homogeneity, distribution, and separation of cell types by clustering in UMAP plots. Minority cell types in a cluster, defined as cell type that accounts for less than 5% of the total counts for that cluster, were considered as potential hybrid cells and excluded from further analysis.

## Cell abundance analysis

To determine whether the risk factors of interest- age, sex, and genotype-contribute to the variance in cell type abundance, we compared cell fractions per biological sample within each cluster across different sample groups. Briefly, the fraction calculation involved dividing the cell count of a biological sample in a specific cluster by the total cell counts for that sample across the entire dataset.

Within each cluster, we grouped biological replicates (n = 3 or 4) and compared the distribution of these fractions across various sample groups using a three-way-ANOVA test. We employed the three-way-ANOVA test to investigate whether the independent variables (the three risk factors) or their interactive effects significantly contribute to the observed variation in outcomes, specifically, the difference in cell abundance. This analysis was conducted using the *aov* function within the Stats package (version 3.6.2) in R.

## Cell cycle scoring and annotation

To assess the variation in the distribution of cell phases within a specific cell type across different sample groups, we estimated the cell cycle phase of each cell. This was accomplished by calculating scores for each phase using the *CellCycleScoring* function in Seurat. This function predicts the classification of each cell into G1, G2M, or S phases based on lists of cell cycle markers obtained from Tirosh et al, as explained in the Seurat tutorial. Subsequently, cells were labeled with the cycle phase corresponding to the highest score. We computed the proportion of each cell cycle phase within a specific cell type and compared the proportions of each phase across sample groups. Finally, we applied a three-way-ANOVA test to determine whether the risk factors under consideration-age, sex, or genotype - significantly contributed to the observed variance in cell phase abundance across sample groups.

## Cell-type-specific differential expression analysis

For each cell type, we performed a differential gene expression analysis comparing *APOE4/4* cells to *APOE3/3* cells and stratified the analysis by age and sex groups. This analysis was conducted using the *FindMarkers* function in the Seurat package. We set the test.use parameter to MAST, a two-part generalized linear model, as recommended by Mou et al. in a study comparing 9 DE methods for single-cell RNA sequencing analysis^50^. MAST models gene expression rate using linear regression and expression level using Gaussian distribution. Differentially expressed genes (DEGs) were considered significant based on Bonferroni-corrected adjusted p-value cutoff of less than 0.05 and a log2 fold change (LFC) value greater than 0.1.

## Pathway enrichment analysis

We performed pathway enrichment analysis using a web tool g:Profiler^51^, which conducts functional over-representation analysis by associating a given gene list with known functional information sources and identifies statistically significant enriched terms. For each cell-type-specific DEG list, we separated the upregulated and downregulated subsets and independently queried them with an adjusted p-value cutoff of 0.05 to identify significant pathway enrichments in both directions. We employed the g:SCS algorithm^52^, a default method for multiple testing correction in pathway enrichment analysis, to account for dependencies among hierarchically related pathway terms. Our analysis encompassed terms from Gene Ontology cellular components, biological processes, and molecular functions, as well as Human Protein Atlas, Human Phenotype Ontology, KEGG, Reactome, and Wiki pathways.

## Enriched pathway network visualization

Following a previously established protocol, we conducted network pathway analysis on the pathway results derived from our cell-type-specific DEGs. In summary, the cell-type-specific upregulated and downregulated pathways were imported into the Cytoscape^53^ visualization application, EnrichmentMap^54^, individually, and distinguished by color intensities. Upregulated pathways were labeled with dark colors, while downregulated pathways were labeled with light colors. Each node, representing a pathway, was filled with different color tones to indicate their cell type associations. We marked nodes green for excitatory neurons, blue for inhibitory neurons, yellow for astrocytes, red for microglia, purple for oligodendrocytes, and pink for oligodendrocyte precursor cells. Redundant and related pathways were consolidated into single biological themes using AutoAnnotate, a Cytoscape application. Pathway nodes are connected by edges if the two pathways share gene set overlaps. Labels for biological-themed clusters were generated with the WordCloud algorithm using the “Adjacent Words” option.

## Cell-cell communication

Cell-cell communication (CCC) was calculated using the CellChat^55^ method. Briefly, CellChat utilizes ligand, receptor, and cofactor expression from transcriptomic data to calculate a CCC probability. First, based on a CellChat-curated database of ligand-receptor interactions, differentially expressed signaling genes are used to calculate ensemble average expression of signaling genes. Communication probability is modeled using the law of mass action, and statistically significant communications are identified using a permutation test. We then evaluated the interaction across cell types (excitatory neurons, inhibitory neurons, oligodendrocytes, oligodendrocyte precursor cells, and astrocytes).

## Quantification of gene expression metrics

Single-cell expression values for the *hApoE* transgene were extracted from the Seurat object using the SCT-normalized assay. Briefly, we set the default assay to SCT and obtained per-cell expression values for the gene of interest from the "data" slot, which contains the Pearson residuals returned by SCTransform. For each cell, we also retrieved its corresponding experimental group label from the combined metadata variable age_sex_geno, which encodes age, sex, and APOE genotype.

To summarize gene expression at the group level, we first defined “positive” cells as those with SCT residuals strictly greater than zero. For each group, we then computed: (i) the total number of cells with non-missing expression (n_total), (ii) the number of positive cells (n_pos), (iii) the percentage of positive cells (pct_pos = 100 × n_pos / n_total), (iv) the mean expression across all cells in the group (avg_all), and (v) the mean expression restricted to positive cells only (avg_pos), defined as the mean of expression among cells with residuals > 0. Groups with no positive cells were assigned NA for avg_pos. All summaries were implemented in R using the dplyr package.

## Statistical comparisons across groups for APOE expression

To assess pairwise differences between groups, we performed three sets of pairwise comparisons: (1) average expression across all cells (avg_all), (2) average expression among positive cells only (avg_pos), and (3) the fraction of positive cells (percent_pos = n_pos / n_total). For avg_all, we used a non-parametric Wilcoxon rank-sum test (Mann–Whitney U) on the per-cell expression values, comparing every pair of groups and adjusting the resulting p-values for multiple testing using the Benjamini–Hochberg (BH) method. For avg_pos, we repeated the same Wilcoxon procedure but restricted the analysis to cells classified as positive (expression > 0); pairwise tests were only carried out for group pairs with at least one positive cell in each group. For the fraction of positive cells, we used pairwise tests for equality of binomial proportions (R function pairwise.prop.test), again with BH correction for multiple comparisons.

---

**Extended Data Fig. 1.**
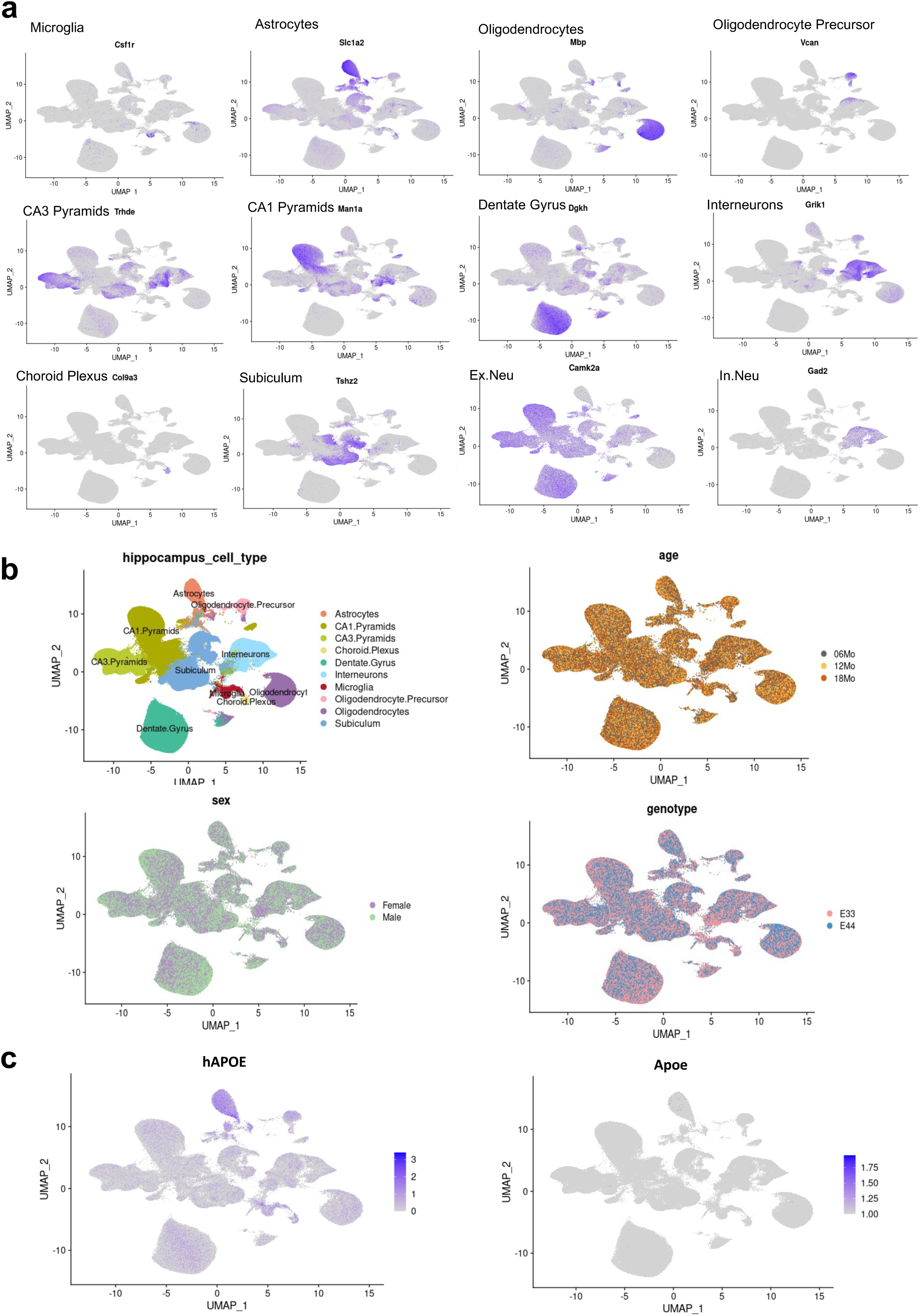
| Characterization of cell identities, neuronal subpopulations, and *APOE* expression patterns. **a.** Feature plots of canonical marker genes for major brain cell types, illustrating expression patterns that guide cell type identification. **b.** UMAP embeddings annotated by neuronal subpopulations, showing distinctions defined by hippocampal regional identity, and by risk factors: age, sex, and *APOE* genotype. **c.** UMAP embeddings highlighting the expressions of the human *APOE* transgene and endogenous mouse *Apoe*, shown with enhanced color contrast to visualize their spatial patterns across cell types. Scale indicates expression levels.

**Extended Data Fig. 2.**
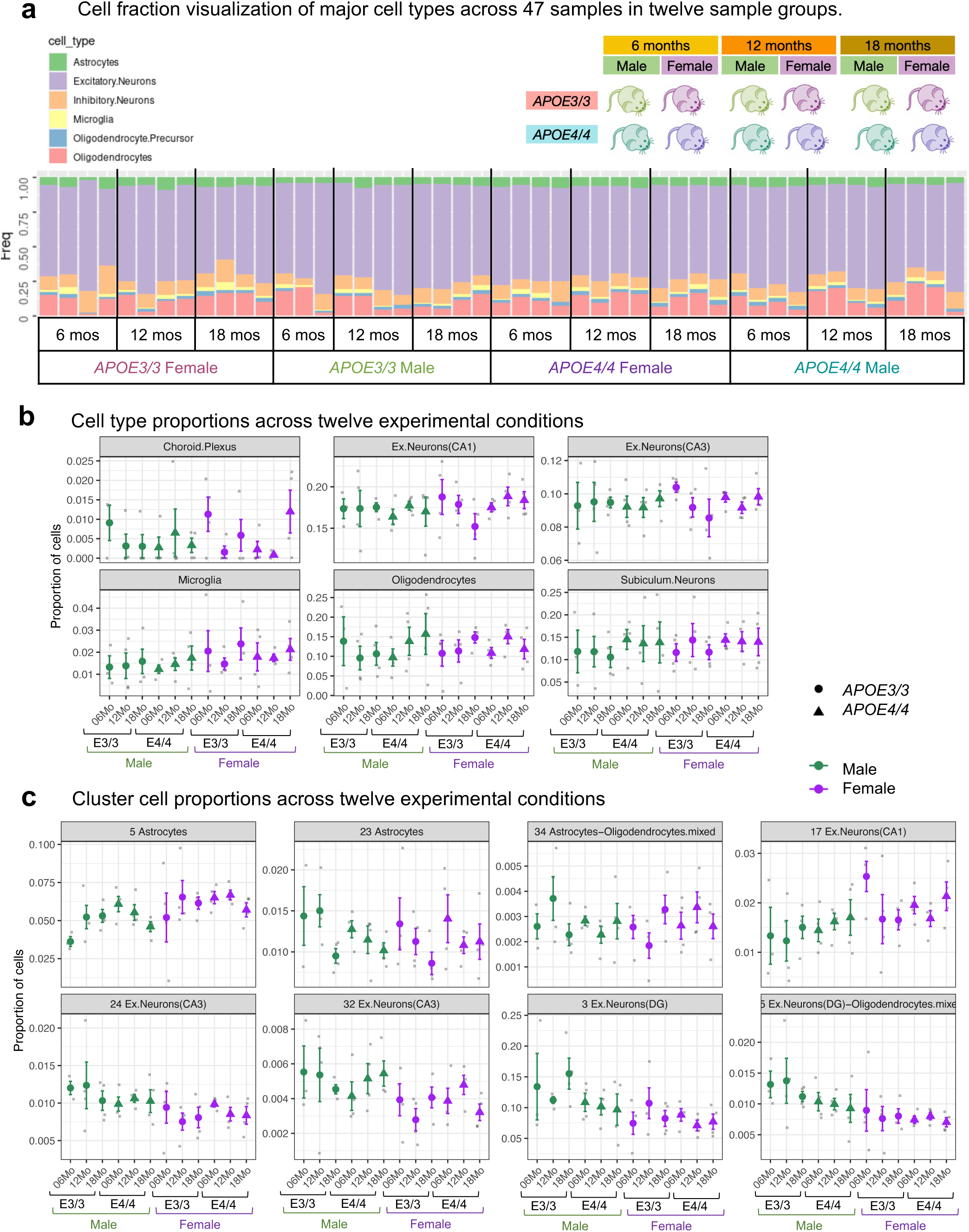
| Cell type and subcluster distribution across the twelve experimental conditions. **a.** Cell type composition for each sample and across all twelve experimental conditions. **b.** Cell type proportions across the twelve conditions for choroid plexus, CA1 and CA3 excitatory neurons, microglia, oligodendrocytes, and subiculum neurons. **c.** Subcluster-level cell proportions per sample across the twelve conditions for subclusters that showed significant effects in the three-way ANOVA analysis.

**Extended Data Fig. 3.**
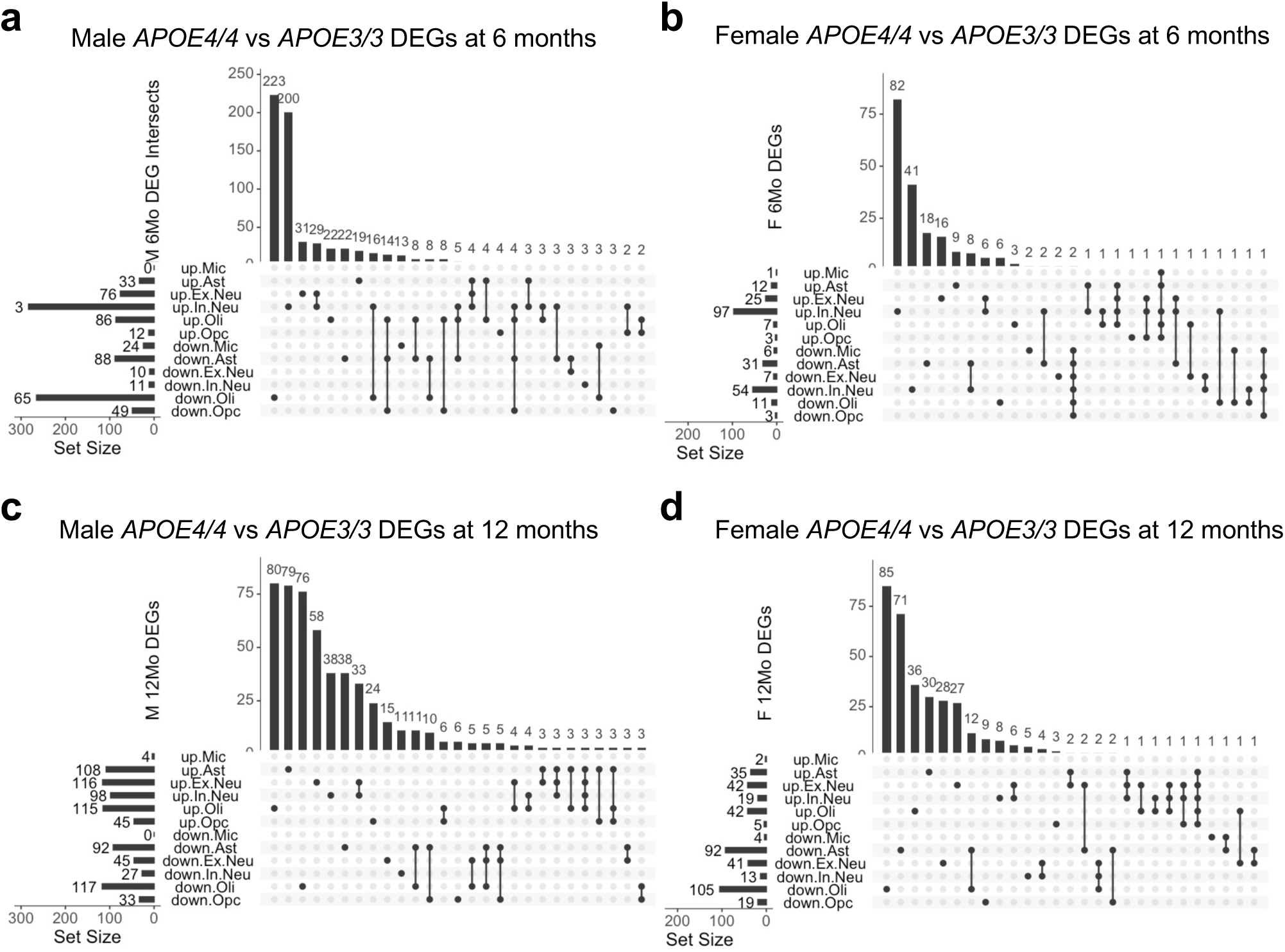
| Differentially expressed gene (DEGs) intersections across cell types at 6 and 12 months. a–d. UpSet plots showing DEG overlaps (*APOE4/4* vs. *APOE3/3*) among the six major cell types in males (**a,c**) and females (**b,d**) at 6 months (**a,b**) and 12 months (**c,d**).

**Extended Data Fig. 4.**
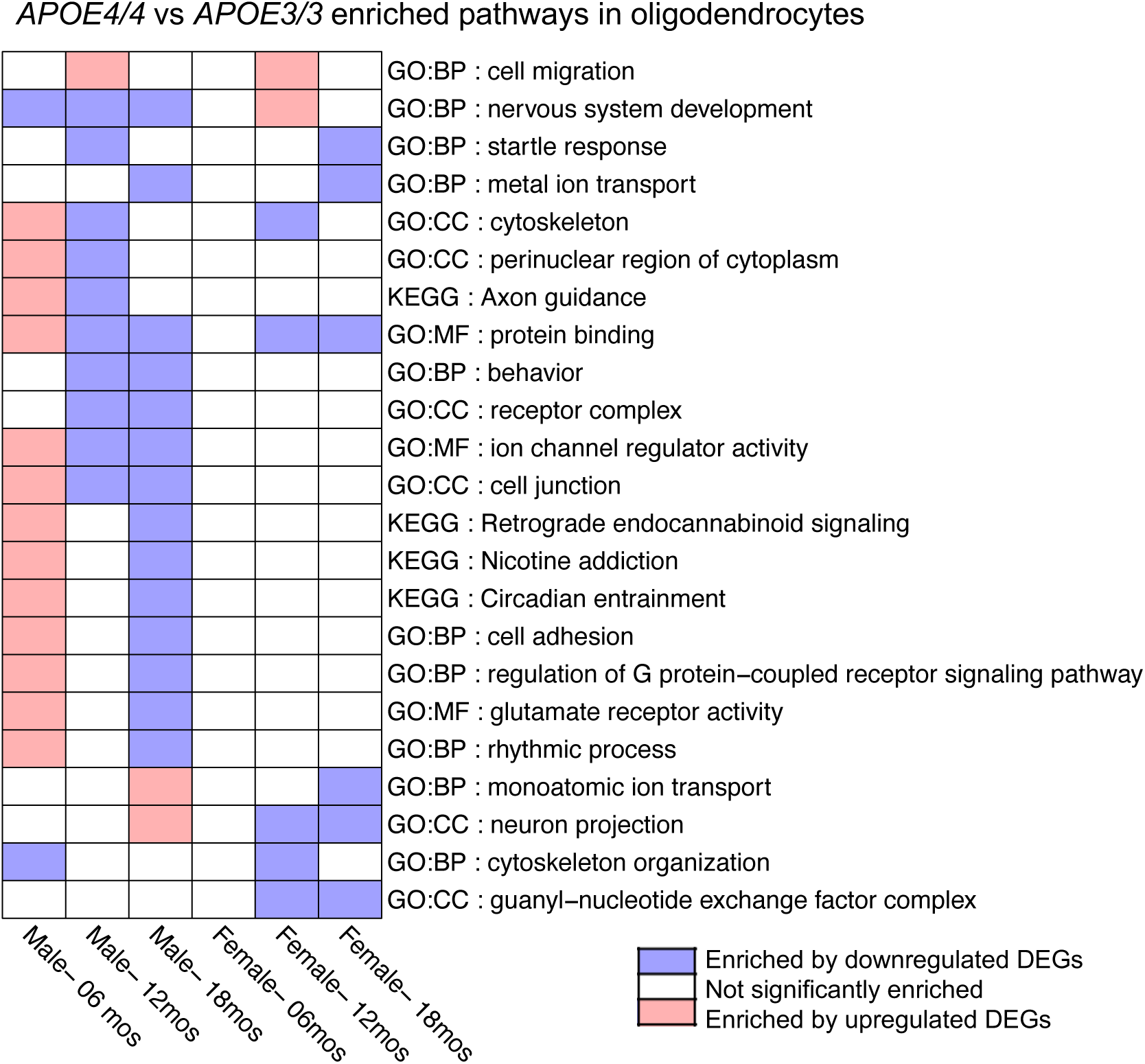
| Heatmap of *APOE4/4* versus *APOE3/3* enriched pathways in oligodendrocytes, stratified by age and sex. Colors indicate the direction of regulation, with red denoting upregulation, blue denoting downregulation, and white indicating non-significant changes in the corresponding groups.

**Extended Data Fig. 5.**
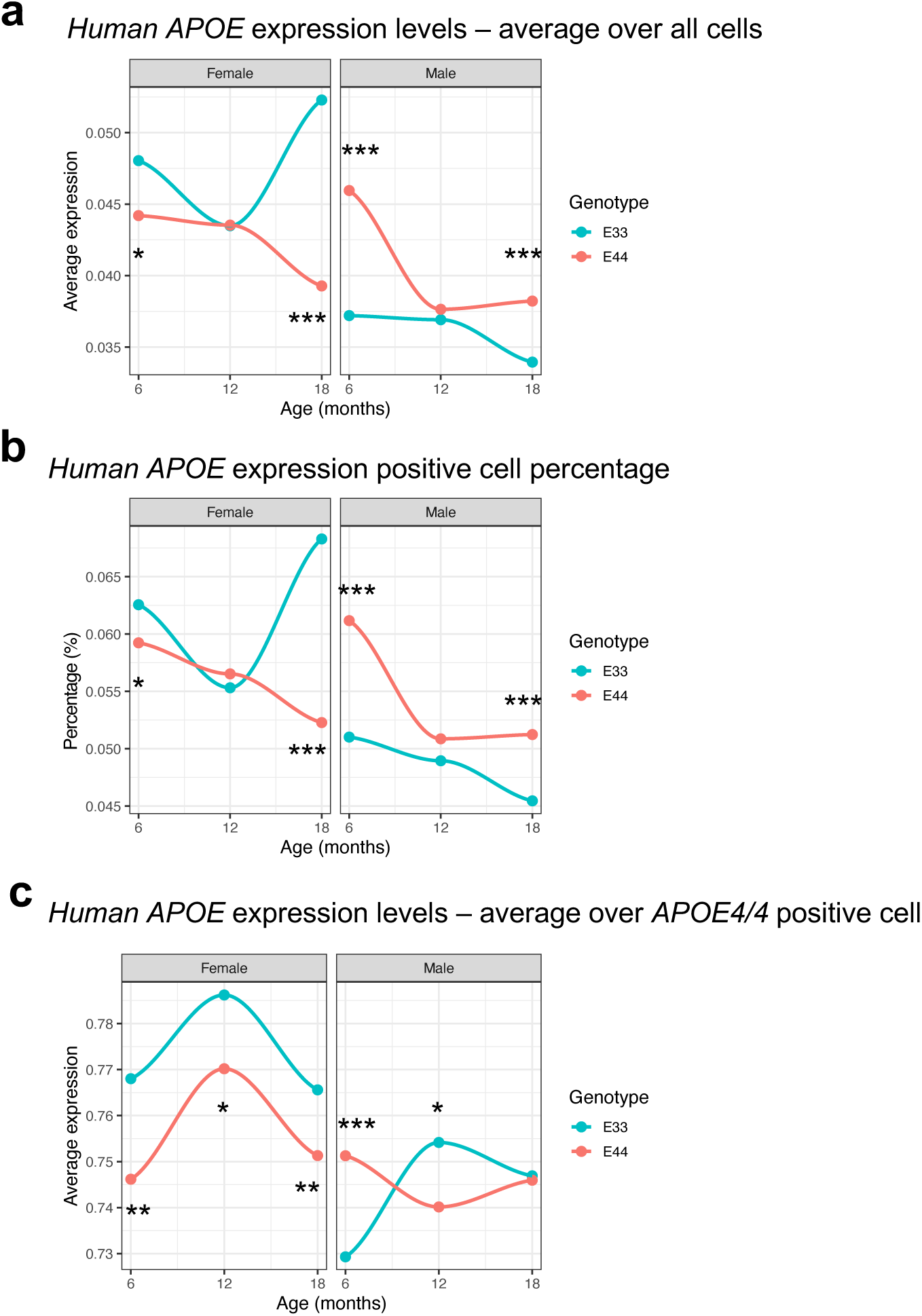
| *APOE* expression levels comparison between *APOE* genotypes in females and males across ages. a. Average human *APOE* expression across all cells within each sample. b. Percentage of *APOE* expression-positive cells within each sample. c. Average human *APOE* expressions normalized only among *APOE* expression-positive cells within each sample. a-c. Significance levels for *APOE4*/4 versus *APOE3*/3 comparisons are indicated as follows: * for p < 0.05, ** for p < 0.01, and *** for p < 0.001.

**Extended Data Fig. 6.**
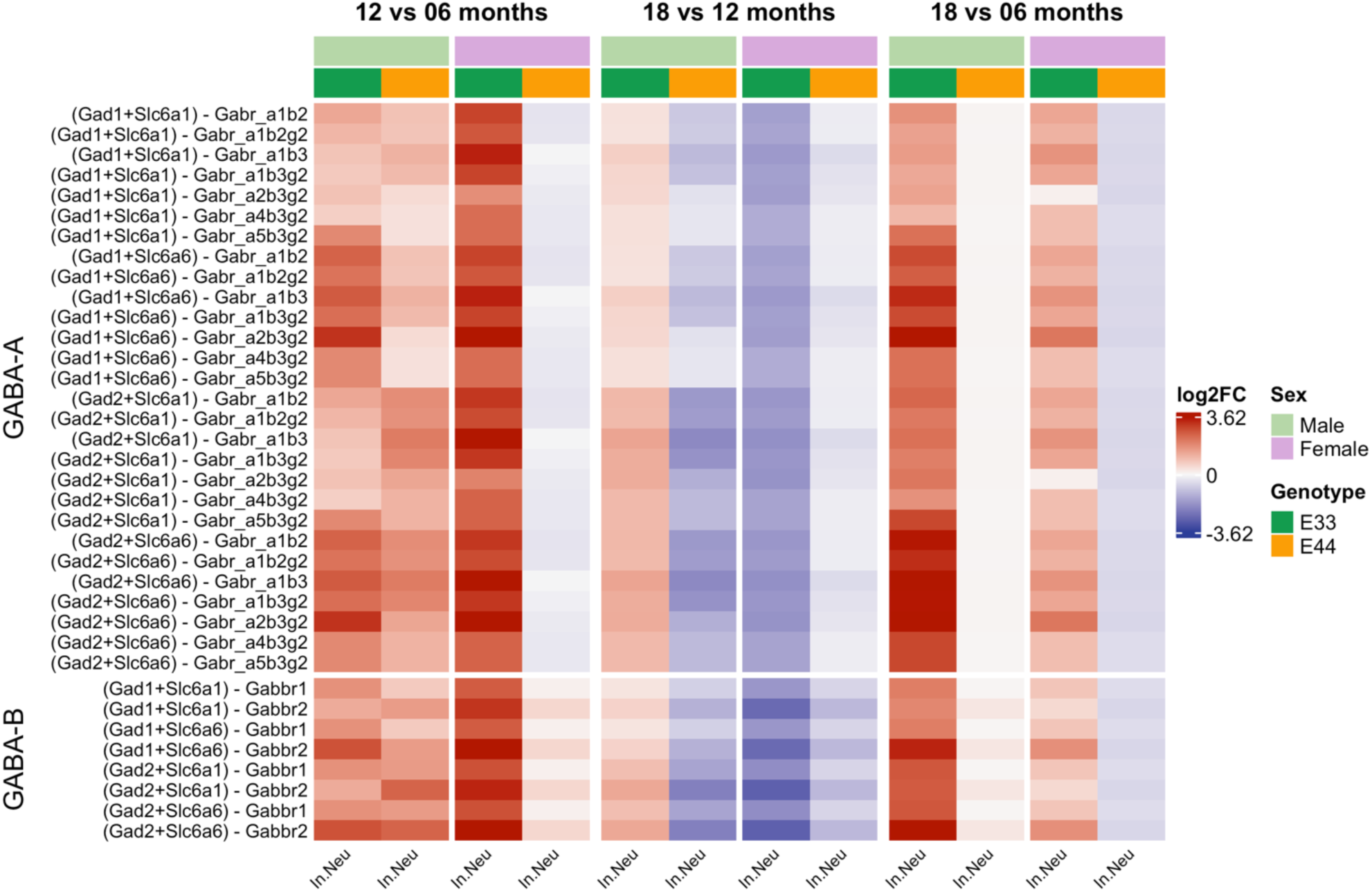
| Heatmap of cell communication probability log2 fold-change of older age compared to younger age of GABA-A and GABA-B ligand-receptor pairs in inhibitory neurons. Ligands are represented in parentheses, followed by the corresponding receptor.

